# Knowledge syntheses in medical education: A bibliometric analysis

**DOI:** 10.1101/2020.05.12.088542

**Authors:** Lauren A. Maggio, Joseph A. Costello, Candace Norton, Erik W. Driessen, Anthony R. Artino

## Abstract

**Purpose:** This bibliometric analysis maps the landscape of knowledge syntheses in medical education. It provides scholars with a roadmap for understanding where the field has been and where it might go in the future. In particular, this analysis details the venues in which knowledge syntheses are published, the types of syntheses conducted, citation rates they produce, and altmetric attention they garner.

**Method:** In 2020, the authors conducted a bibliometric analysis of knowledge syntheses published in 14 core medical education journals from 1999 to 2019. To characterize the studies, metadata was extracted from Pubmed, Web of Science, Altmetrics Explorer, and Unpaywall.

**Results:** The authors analyzed 963 knowledge syntheses representing 3.1% of total articles published (n=30,597). On average, 45.9 knowledge syntheses were published annually (SD=35.85, Median=33), and there was an overall 2,620% increase in the number of knowledge syntheses published from 1999 to 2019. The journals each published, on average, a total of 68.8 knowledge syntheses (SD=67.2, Median=41) with *Medical Education* publishing the most (n=189; 19%). Twenty-one knowledge synthesis types were identified; the most prevalent types were systematic reviews (n=341; 35.4%) and scoping reviews (n=88; 9.1%). Knowledge syntheses were cited an average of 53.80 times (SD=107.12, Median=19) and received a mean Altmetric Attention Score of 14.12 (SD=37.59, Median=6).

**Conclusions:** There has been considerable growth in knowledge syntheses in medical education over the past 20 years, contributing to medical education’s evidence base. Beyond this increase in volume, researchers have introduced methodological diversity in these publications, and the community has taken to social media to share knowledge syntheses. Implications for the field, including the impact of synthesis types and their relationship to knowledge translation, are discussed.

## Introduction

> “There is a need to move from opinion-based education to evidence-based education.”
>
> — - Harden (2000)^1^

As the 20th century came to a close, Harden challenged the field of medical education to develop its evidence base in an effort to empower medical educators to act as evidence-informed teachers.^1^ In response to this call and inspired by the evidence-based medicine movement, the Association for Medical Education in Europe established the Best Evidence in Medical Education (BEME) Consortium, which called upon the medical education research community to create knowledge syntheses. More than two decades later, medical education has seen a steady rise in knowledge syntheses; yet, there has been limited effort to understand this growth and map the landscape of these knowledge syntheses in medical education. This lack of understanding is problematic because without knowing where the field has been it is difficult to chart our future to ensure a robust evidence base moving forward.

Knowledge syntheses are defined as “the contextualization and integration of research findings of individual research studies within the larger body of knowledge on the topic”.^3^ Gordon asserts that in medical education the creation of knowledge syntheses is as “vital as the production of novel primary research”.^2^ Knowledge syntheses can provide medical educators with the means to move from reliance on eminence—and experience-based opinion or individual studies—to the utilization of resources that holistically consider and integrate the best available evidence.^4^ Thus, it is unsurprising that interest in knowledge syntheses has resulted in the publication of several instructional guides on how to conduct knowledge syntheses in medical education^5–7^ and the establishment of grant funding opportunities to support their creation.^8^ What is more, systematic reviews are often a gateway for medical students starting their own lines of research^5^ and are also essential for graduate students entering the field. To this end, BEME reports a 50% increase in the number of reviews undertaken by authors over a recent five-year period.^9^

Over the last decade, broad bibliometric reviews of the literature have been undertaken to identify trends in medical education publishing,^10–12^ recognize the most cited articles in the field,^13^ characterize author networks across international borders,^14^ and describe associated social media attention.^15,16^ While these studies provide important insights into the overall landscape of the medical education literature, they fail to fully illuminate knowledge syntheses and their characteristics. More recently, members of our author team conducted a meta-synthesis of BEME knowledge syntheses, attempting to identify the degree to which the reviews were ready for translation into practice and to summarize their features.^17^ Although valuable, this metasynthesis examined a limited sample of knowledge syntheses (BEME reviews only) that underwent several intense review steps prior to publication; as such, the generalizability of this work is limited.

The bibliometric analysis reported here aims to map the broader landscape of knowledge syntheses in medical education to provide scholars with a roadmap for understanding both where the field has been and where it might go in the future. In particular, we examine the venues in which knowledge syntheses are published, the types of syntheses conducted, the citation rates they produce, and the altmetric attention they garner. In doing so, we hope to provide a detailed summary of the characteristics of medical education knowledge syntheses over the past 20 years.

## Methods

We conducted a bibliometric analysis of knowledge syntheses. A bibliometric analysis aims to use quantitative methods to describe characteristics of publications (e.g., journal articles) and their publication patterns, in order assess the current state of a field and provide insight into its overall structure.^18^ Although we did not conduct a systematic review or meta-analysis, where appropriate, we report our methods in alignment with the Preferred Reporting Items for Systematic Reviews and Meta-Analyses^19^ to provide transparency and facilitate replication of our approach.

### Data Collection

On March 26, 2020 we queried PubMed using a search strategy designed by LM and JC, two researchers trained in information science. The purpose of this systematic search was to broadly capture knowledge syntheses using a combination of keywords, such as knowledge synthesis, literature review, meta-synthesis, and controlled vocabulary terms (Appendix 1). We limited our search to the last 20 years (1999 to 2019) to correspond with the establishment of the BEME initiative.

We restricted our search to 14 journals that have been previously identified as core medical education journals.^10,17^ The journals included in our analysis were: *Academic Medicine, Advances in Health Sciences Education, BMC Medical Education, Canadian Medical Education Journal, Clinical Teacher, International Journal of Medical Education, Advances in Medical Education and Practice, Journal of Graduate Medical Education, Medical Education, Medical Education Online, Medical Teacher, Perspectives on Medical Education, Teaching and Learning in Medicine*, and *The Journal of Continuing Education in the Health Professions*. We downloaded the metadata for all resulting citations from PubMed into GoogleSheets. We also searched PubMed for all articles published in these journals over the study period. This enabled us to contextualize knowledge syntheses within the broader scope of the literature.

While all of these journals are indexed in PubMed, seven of them are not included in their entirety, including *Advances in Health Sciences Education, Canadian Medical Education Journal, Clinical Teacher, Medical Education Online, Medical Teacher, Teaching and Learning in Medicine*, and *The Journal of Continuing Education in the Health Professions*. For example, *Clinical Teacher* first appeared in PubMed in 2010, but the journal started publishing articles in 2003. Therefore, for these seven journals we identified citations from these periods by hand searching the journal’s website or Web of Science (WoS). This supplemental searching was conducted to identify both knowledge syntheses as well as all other articles published in the journal sample.

On March 31, 2020 we queried Altmetric Explorer to obtain information on each article’s Altmetric Attention Score, which is a web-based measure of an article’s impact, with an emphasis on social media outlets as sources of data (e.g., whether the article was tweeted, saved to Facebook, covered by the news media, etc.). Next, on April 10, 2020 we searched WoS to obtain additional metadata (e.g., number of times cited, funding information). To provide context in the broader field of medical education, we also downloaded citation and Altmetric data for all articles in the sample for the selected time period. Lastly, on March 31, 2020, we queried the UnPaywall database, which tracks the accessibility of journal articles, to determine each article’s open access status. All data were downloaded and managed in GoogleSheets (Google, 2020).

### Inclusion Criteria

Knowledge syntheses were included if they met the above definition as written by Canadian Institute of Health Research.^3^ We also included knowledge syntheses in which authors reviewed and synthesized the content and structure of research studies to contribute understanding on a topic (e.g., knowledge syntheses that defined a concept based on the literature or identified how studies were conducted). We excluded articles focused solely on the bibliometric properties of knowledge and articles focused on the mechanics of conducting a knowledge synthesis. We also excluded articles that solely analyzed documents that were not findings of research studies, such as syntheses of promotion and tenure guidelines or curricular documents. We also excluded articles that included a review of the literature, but were primarily focused on powering an additional research approach such as a Delphi study or survey.

To determine inclusion, LM and JC independently reviewed the titles and abstracts of all citations and then met via conference call to discuss coding discrepancies. TA was available to facilitate any coding disagreements.

### Data extraction

All data was extracted from each included article’s citation metadata. For example, to determine review type (e.g., systematic review, scoping review, etc.), we relied solely on the text available in the title or abstract.

### Analysis

Descriptive statistics were calculated using GoogleSheets^20^ and IBM SPSS Statistics^21^. We also calculated a Spearman’s rank-order correlation to examine the relationship between the age of the knowledge syntheses and two of the collected variables.

## Results

We identified 2,210 citations of which 963 met inclusion criteria (Fig. 1). Included citations represented 3.1% of total articles published in the overall journal sample between 1999 and 2019 (n=30,597).

**Figure 1:**
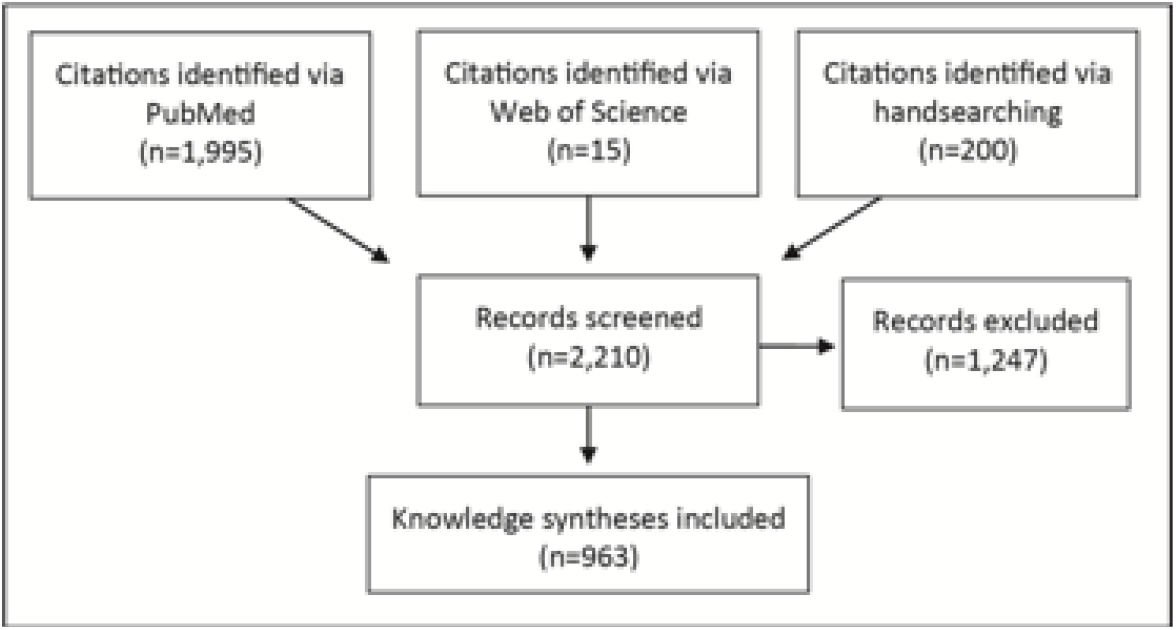
Knowledge synthesis identification

Knowledge syntheses were published in all years studied (1999 to 2009) with 49% (n=470) published between 2015 and 2019, and 84% (n=805) published between 2009 and 2019 (Fig 2). On average, there were 45.9 knowledge syntheses published annually (SD=35.85, Median=33). The fewest knowledge syntheses were published in 1999 (n=5) and the most in 2019 (n=136), representing an overall percent increase of 2,620% across the 20-year period. In contrast, over the same time period, non-knowledge synthesis articles in these journals experienced an overall percent increase of 204%.

**Figure 2:**
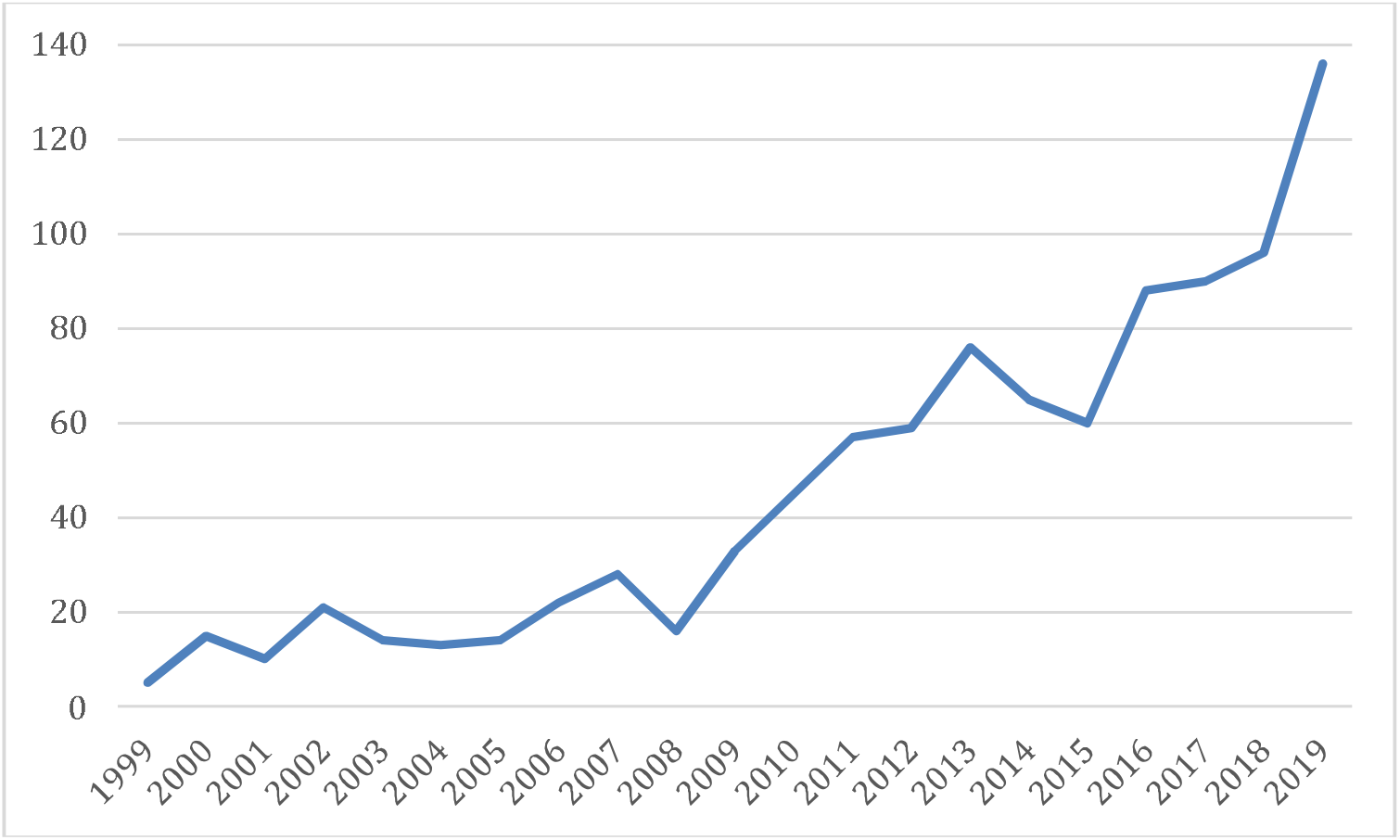
The number of knowledge syntheses published in 14 core medical education journals between 1999 and 2019

### Journal coverage

All journals published at least one knowledge synthesis (Table 1). On average journals published a total of 68.8 knowledge syntheses (SD=67.2, Median=41). *Medical Education* published the most knowledge syntheses overall (n=189; 19%) followed by *Academic Medicine* (n=186; 19%) and *Medical Teacher* (n=172; 18%). The *Canadian Medical Education Journal* published the fewest number of knowledge syntheses (n=10).

**Table 1:** Knowledge Syntheses (KS) by journal

| Journal | Data Available | Total Articles | Total KS (% total) | Top KS type of those specified (n, %) | Non-specified KS type (%) |
| --- | --- | --- | --- | --- | --- |
| *Acad Med* | 1999-2019 | 8450 | 187 (2.2) | Systematic (70, 37.4) | 60 (32.1) |
| *Adv Health Sci Educ Theor Pract* | 1999-2019 | 1142 | 58 (5.1) | *Systematic, Systematic w/ Meta-Analysis (8, 13.8) | 24 (41.4) |
| *Adv Med Educ* | 2010-2019 | 775 | 40 (5.2) | Systematic (10, 25.0) | 17 (42.5) |
| *BMC Med Educ* | 2001-2019 | 2766 | 116 (4.2) | Systematic (59, 50.9) | 18 (15.5) |
| *Can Med Educ J* | 2010-2019 | 318 | 10 (3.1) | Systematic (3, 30.0) | 4 (40.0) |
| *Clin Teach* | 2004-2019 | 1543 | 20 (1.3) | Narrative (4, 20.0) | 13 (65.0) |
| *Int J Med Educ* | 2011-2019 | 337 | 13 (3.9) | *Systematic, Narrative (2, 15.4) | 7 (53.9) |
| *J Cont Educ Health Prof* | 1999-2019 | 884 | 45 (5.1) | Systematic (16, 35.6) | 22 (48.9) |
| *J Grad Med Educ* | 2009-2019 | 1835 | 42 (2.3) | Systematic (16, 38.1) | 10 (23.8) |
| *Med Educ* | 1999-2019 | 5296 | 189 (3.6) | Systematic (58, 30.7) | 70 (37.0) |
| *Med Educ Online* | 1999-2019 | 700 | 20 (2.9) | Systematic (7, 35.0) | 8 (40.0) |
| *Med Teach* | 1999-2019 | 4941 | 172 (3.5) | Systematic (77, 44.8) | 57 (33.1) |
| *Perspect Med Educ* | 2012-2019 | 517 | 22 (4.3) | Systematic (6, 27.3) | 6 (27.3) |
| *Teach Learn Med* | 1999-2019 | 1093 | 29 (2.7) | Systematic (6, 20.7) | 16 (55.2) |

\*Represents a tie

### Knowledge synthesis types

We identified 21 knowledge syntheses types, which accounted for 631 publications. Knowledge synthesis type was unspecified for 34% (n=332) of publications. Systematic reviews were the most prevalent knowledge synthesis type (n=341; 35.4%) followed by scoping reviews (n=88; 9.1%), systematic reviews with meta-analysis (n=62, 6.4%), and narrative reviews (n=50; 5.2%) (Table 2). Additionally, systematic reviews were the top knowledge synthesis type for each journal except for *Clinical Teacher*, which predominantly published narrative reviews. The following review types each represented less than 10 articles (<10% of the sample): qualitative, thematic, historical, guideline, meta-narrative, synthetic, rapid, meta-review, methodological, systematic scoping, umbrella, mapping, meta-synthesis, and meta-ethnography.

**Table 2:** Knowledge syntheses types

| Type | Number (%) | Year first appeared in sample | Average Citations (SD, Median) n=847 | Average Altmetric Attention Score (SD, Median) n=769 |
| --- | --- | --- | --- | --- |
| Systematic | 341 (35.4) | 1999 | 57.6 (134.1, 21) | 16.5 (53.8, 6) |
| Not Specified | 332 (34.4) | 1999 | 66.1 (87.6, 30) | 10.5 (15.6, 5) |
| Scoping | 88 (9.1) | 2011 | 7.8 (12.3, 3) | 13.6 (12.9, 10) |
| Systematic w/ Meta-analysis | 62 (6.4) | 2006 | 52.8 (90.1, 17) | 23.6 (51.6, 7) |
| Narrative | 56 (5.8) | 2006 | 26.1 (39.8, 10) | 8.7 (11.0, 4) |
| Critical | 35 (3.6) | 1999 | 80.4 (149.4, 46) | 10.8 (21.8, 4) |
| Integrative | 12 (1.2) | 2004 | 12.3 (15.5, 9) | 5.9 (3.4, 5) |
| Realist | 11 (1.1) | 2010 | 33.6 (58.7, 11) | 19.6 (9.9, 16) |

### Funding

Funding was reported for 285 (30%) knowledge syntheses, with systematic reviews (n=122; 43%) being the most commonly funded knowledge synthesis type. Forty (15%) of these knowledge syntheses reported more than one funder. The US National Institutes of Health was the most commonly reported funder (n=41; 14%). However, a variety of funders were credited for their support including, but not limited to, national and regional agencies (e.g., the Canadian Institute of Health Research,^22^ the European Commission;^23^) professional societies (e.g., such as the Society of Directors of Research in Medical Education^8,24^ and Society of Hospitalist Pharmacists of Australia;^25^) and foundations (e.g, the Arnold P. Gold Foundation and the Bill and Melinda Gates Foundation^26^). Multiple authors listed funding assistance from their home institutions.

### Citations

Knowledge syntheses, for which we were able to extract data from WoS (n=847), generated a total of 45,566 citations. Citation counts ranged from 0 to 1,629 and, on average, knowledge syntheses were cited 53.80 times (SD=107.12, Median=19). For comparison, over the same time period, all other articles published in these journals (e.g., research reports, editorials, perspectives, commentaries, etc.) were cited an average of 15.30 times (SD=39.22, Median=5).

Fifty-one knowledge syntheses had not been cited at the time of data extraction. Of these uncited knowledge syntheses, 41 (80%) were published in 2019. The most cited knowledge synthesis, a 2005 BEME systematic review on simulation, has received 1,602 citations (Table 3). The top three cited knowledge synthesis types, of those that were specified, were critical reviews (Mean=80.35, SD=149.37, Median=46), systematic reviews (Mean=57.64, SD=134.06, Median=21), and systematic reviews with meta-analysis (Mean=52.76, SD=90.06, Median=17) (Table 2).

**Table 3:** Top 10 Cited Knowledge Syntheses

| Knowledge Syntheses (publication year) | First Author | Citations | KS Type | Freely Accessible | Altmetric Attention Score |
| --- | --- | --- | --- | --- | --- |
| Features and uses of high-fidelity medical simulations that lead to effective learning: a BEME systematic review (2005)<sup>27</sup> | Issenberg, S. | 1629 | Systematic Review | No | 26 |
| Systematic review of depression, anxiety, and other indicators of psychological distress among US and Canadian medical students (2006)<sup>28</sup> | Dyrbye, L. | 964 | Systematic Review | Yes | 65 |
| Reflection and reflective practice in health professions education: a systematic review (2009)<sup>29</sup> | Mann, K. | 771 | Systematic Review | No | 27 |
| Standards for Reporting Qualitative Research: A Synthesis of Recommendations (2014)<sup>24</sup> | O'Brien, B. | 711 | Guideline | Yes | 86 |
| A critical review of simulation-based medical education research: 2003-2009 (2010)<sup>30</sup> | McGaghie, W. | 699 | Critical Review | No | 37 |
| A systematic review of faculty development initiatives designed to improve teaching effectiveness in medical education: BEME Guide No. 8 (2006)<sup>31</sup> | Steinert, Y. | 634 | Systematic Review | No | 21 |
| Toward a definition of competency-based education in medicine: a systematic review of published definitions (2010)<sup>32</sup> | Frank, J. | 613 | Systematic Review | No | 20 |
| Does Simulation-Based Medical Education With Deliberate Practice Yield Better Results Than Traditional Clinical Education? A Meta-Analytic Comparative Review of the Evidence (2011)<sup>33</sup> | McGaghie, W. | 571 | Systematic Review w/ Meta-analysis | Yes | 40 |
| Effectiveness of problem-based learning curricula: Research and theory (2000)<sup>34</sup> | Colliver, J. | 533 | Critical Review | Yes | 0 |
| A best evidence systematic review of interprofessional education: BEME Guide no. 9 (2007)<sup>35</sup> | Hammick, M. | 520 | Systematic Review | No | 6 |

There was a strong positive correlation between the age of the synthesis and its citation rate (Spearman’s □=.79, p<.01), indicating that older reviews were cited more often than more recent reviews.

### Altmetrics

The majority of knowledge syntheses received social media attention (i.e., altmetric attention; n=771; 80%) generating 15,149 total mentions across 10 altmetric outlets (e.g., Twitter, Facebook). Knowledge syntheses were collectively saved by 92,598 Mendeley readers. Articles in all journals except for the *Journal of Continuing Education in the Health Professions* received social media mentions. Altmetric Attention scores ranged from 0 to 806; Mean=14.12 (SD=37.59, Median=6). For comparison, over the same time period, all other articles published in these journals received an average Altmetric Attention score of 7.46 (SD=32.21, Median=3).

There was a weak negative correlation between the age of the synthesis and its Altmetric Attention score (Spearman’s □=-.24, p<.01), indicating that newer reviews attained a slightly higher altmetric score than older reviews.

The majority of social media attention was received on Twitter (n=14,172 tweets), with activity generated by 6,308 unique tweeters from 105 countries. The next most prevalent sources of altmetric attention were Facebook (n=314 posts), the news media (n=251 mentions), blogs (n=200 posts), and policy documents (n=99) (Table 4). Articles were not mentioned in seven additional outlets tracked by Altmetric, including patent documents, Pinterest, Weibo, LinkedIn, Q&A resources, Reddit, and syllabi.

**Table 4:** Altmetric mentions and Mendeley readers of knowledge syntheses (KS) published between 1999 and 2019.

| Mention type | Total Mentions for Knowledge Syntheses | KS receiving mention type (%) n=769 |
| --- | --- | --- |
| Twitter | 14,172 | 686 (89.21) |
| Facebook | 314 | 214 (27.83) |
| News Media | 251 | 52 (6.76) |
| Blogs | 200 | 112 (14.56) |
| Policy | 99 | 69 (8.97) |
| Google+ | 54 | 42 (5.46) |
| Wikipedia | 23 | 21(2.73) |
| Video | 6 | 6 (.78) |
| F1000 | 1 | 1 (.13) |
| Peer Review | 1 | 1 (.13) |
| Mendeley Readers | 92,598 | 756 (98.31) |

The article with the highest altmetric score (806) was, “A systematic review of the effectiveness of flipped classrooms in medical education”, published in 2017 in *Medical Education.^36^* This article was mentioned in 87 news media stories. This media coverage focused on the transition from in-person to online learning in the wake of the COVID-19 pandemic. For context, this knowledge synthesis has the second highest altmetric score of all articles published in the selected journals. The next highest knowledge synthesis had an altmetric score of 281.^37^

### Accessibility

Fifty-four percent of knowledge syntheses (n=524) were freely accessible such that a reader could access the full-text of an article without a subscription or without having to pay an access fee. Accessible copies of articles were located on publisher websites and open access repositories (e.g., PubMed Central), including those maintained by the author’s institution. For example, Rees and colleagues deposited to Keele University’s institutional repository the MS Word version of their BEME review: Evidence regarding the utility of multiple mini-interview (MMI) for selection to undergraduate health programmes: a BEME systematic review.^38^ There were five open-access journals in our sample: *Advances in Medical Education and Practice, BMC Medical Education, Canadian Medical Education Journal, Medical Education Online, Perspectives on Medical Education*. Additionally, journals in our sample had varied access policies. For example, *Academic Medicine* makes articles freely available on its website one year after the article appears in print.

## Discussion

Over the past two decades, there has been considerable growth in knowledge syntheses in medical education, including an increased variety of synthesis types and the emergence of scoping reviews as an important approach for synthesizing the medical education literature.

### Increase in knowledge syntheses

Our findings confirm a robust growth in the number of knowledge syntheses in medical education. While the overall growth of non-review journal articles has also been quite large (204% growth in our sample), the growth of knowledge synthesis has been more than 10 times that number (2,620%). We predict this trend will continue, paralleling the growth of funding opportunities (e.g., SDRME’s annual funding opportunity)^8^, medical education graduate programs,^39^ and related supporting initiatives such as BEME and the community’s broader desire for synthesized evidence to keep current with research in the field and improve educational practice.^17^ Additionally, when compared with other article types, knowledge syntheses are more highly cited, which makes them attractive to both authors and editors. For example, 80% of the top cited articles in *Medical Education* were review articles.^40^ In the current academic climate, the count of an individual’s citations are usually factored into faculty promotion and tenure decisions.^41,42^ Also, from an editor’s perspective, journals are typically judged by their impact factor, a citation-based metric. In both cases, this emphasis on citations may make it difficult for authors and editors to pass up the opportunity to write and publish a review article that stands to accrue up to three times as many citations as another article type.

If our prediction of continued growth proves to be correct, then it is critical that knowledge syntheses are optimized for use. A recent meta-synthesis of BEME reviews indicated that while such reviews are quite valuable, there was much room for improvement in relation to accessibility and relevance.^43^ Additionally, a survey of medical educators reported that 20% of participants never or rarely used knowledge syntheses to inform their educational decisions.^44^ From our perspective, this is a lost opportunity for individuals and institutions to advance their evidence-informed practice. In many ways, it is also a waste of resources if one considers the substantial time and energy required to complete a high-quality knowledge synthesis (that might never get translated into practice).^7^ Future research should consider identifying potential barriers to uptake of knowledge syntheses specific to medical education and attempt to identify best practices for their creation, dissemination, and ultimate translation into practice.

### Knowledge syntheses types

We identified 21 knowledge syntheses types. This variety aligns with research in other fields that have identified the presence of between 14 and 25 different knowledge syntheses types.^45,46^ The large variety of knowledge synthesis types in medical education appears to be a somewhat recent trend, with only systematic and critical reviews dating back to 1999, and with scoping and realist reviews appearing in just the last decade. Gordon suggests that this proliferation of synthesis types relates to the shifting scope of investigators’ research questions and aims.^2^ He notes that investigators are not only focused on synthesizing literature to answer whether an educational approach works (e.g., by undertaking a systematic review), but also to discern what works, for whom, and in what contexts (e.g., by publishing a realist review) and to understand why it works and how (e.g., by conducting a meta-ethnography).

This increasing variety of knowledge syntheses highlights a need to ensure that consumers of these reviews are aware of the various types in order to understand in which situations they are best applied, as well as to be better able to judge their quality. For example, educators who are most familiar with the positivist tradition of systematic reviews may find themselves in strange territory when reading or attempting to execute a scoping review which, by design, is not designed to provide a “single answer” like a systematic review. Instead, scoping reviews provide readers with a map of the information landscape on a topic in an effort to help readers subjectively interpret the available evidence.^47^ As Thomas et al. recently noted, “the philosophical stance scholars adopt during the execution of a scoping review, including the meaning they attribute to fundamental concepts such as *knowledge* and *evidence*, influences how they gather, analyze, and interpret information obtained from a heterogeneous body of literature.”^44^ Therefore, anyone considering a knowledge synthesis must first understand the various types so that they can then identify the most appropriate synthesis type to answer their research question or satisfy their study aim.^44^ To meet this need, several published articles compare and contrast knowledge syntheses types and provide practical advice for their conduct and use.^45–48^

### Dissemination of knowledge syntheses

Knowledge translation is defined as a dynamic and iterative process that includes the synthesis, dissemination, exchange, and ethically sound application of knowledge to improve health, provide more effective health services and products, and strengthen the health care system.^3^ In this review, we focus on the knowledge synthesis component of the knowledge translation process, which is critical. We propose, however, that increased accessibility for these articles, as indicated by higher altmetric scores for newer reviews, provides a glimpse into efforts to disseminate this knowledge and build a bridge to span the knowledge to practice gap. In this study, we observed that 80% of knowledge syntheses received altmetric attention with mentions predominantly on Twitter. In medical education, tweets have been observed to have a small positive effect on article page views of the tweeted articles.^50^ In addition, we found that 69 knowledge syntheses were integrated into 99 policy documents, which suggests a translation of knowledge into practice. While these findings are promising, we encourage medical education researchers to consider other ways in which social media and policy integration can be leveraged as a knowledge translation tool for sharing knowledge syntheses.

### Limitations

This study should be considered in light of its limitations. In particular, although we conducted a comprehensive search for reviews, it is possible we inadvertently excluded one or more reviews. In addition, our search was limited to 14 core medical education journals, which did not include specialty-specific medical education journals (e.g., *Academic Emergency Medicine)* or medical education journals from specific world regions (e.g., *African Journal of Health Professions Education*). In addition, for citation data, we relied on WoS, which uses selective indexing, and so we were unable to obtain citation data for 125 (13%) knowledge syntheses. Future work might consider alternate citation sources such as Google Scholar, Dimensions, or an amalgamation of such services, with the caveat that each service has a slightly different approach for data collection, each of which has its strengths and weaknesses.

## Conclusion

Over the past 20 years, medical education researchers have begun to answer Harden’s call to develop the field’s evidence base^1^ by publishing nearly a thousand knowledge syntheses across 14 core journals. In addition to a 2,620% increase in knowledge synthesis volume, medical educators have responded by introducing the field to new review types and by utilizing methods, such as posting to social media, for communicating about and sharing knowledge syntheses. As we look to the future of medical education theory, research, and practice, we envision continued production of knowledge syntheses, which will empower our community to act as evidence-informed educators.

## Ethical Approval

Reported as not applicable

## Data

None reported

## Disclaimer

The views expressed in this article are those of the authors and do not necessarily reflect the official policy or position of the Uniformed Services University of the Health Sciences, the Department of Defense, or the U.S. Government.

## Appendix: Search Strings

### PubMed Search String (LIMIT: 1999/01/01 – 2019/12/31)

(“knowledge synthesis” [title/abstract] OR “literature review”[title/abstract] OR “evidence synthesis” [title/abstract] OR “systematic review”[title/abstract] OR review[title] OR “meta-analysis” [Publication Type] OR review [Publication Type] OR “systematic”[sb] OR scoping[title/abstract] OR “meta-synthesis”[title/abstract] OR “narrative review” [title/abstract] OR “critical review”[title/abstract] OR “critical synthesis”[title/abstract] OR “integrative review” [title/abstract] OR “integrative synthesis”[title/abstract] OR “qualitative review”[title/abstract] OR “metastudy”[title/abstract] OR “realist review”[title/abstract] OR “rapid review”[title/abstract] OR “umbrella review” [title/abstract] OR “BEME” [title/abstract] OR “consensus conference”[title/abstract] OR “medline” [title/abstract] OR “cinahl”[title/abstract] OR “PubMed”[title/abstract] OR “embase”[title/abstract] OR “psycInfo”[title/abstract]) AND (“Acad Med”[journal] OR “Adv Health Sci Educ Theory Pract”[journal] OR “Adv Med Educ Pract”[journal] OR “BMC Med Educ”[journal] OR “Can Med Educ J”[journal] OR “Clin Teach”[journal] OR “J Contin Educ Health Prof”[journal] OR “Teach Learn Med”[journal] OR “Perspect Med Educ”[journal] OR “Med Educ”[journal] OR “Med Educ Online”[journal] OR “Med Teach”[journal] OR “J Grad Med Educ”[journal] OR “Int J Med Educ” [journal])

### PubMed Search String for All Citations (LIMIT: 1999/01/01 – 2019/12/31)

(“Acad Med”[journal] OR “Adv Health Sci Educ Theory Pract”[journal] OR “Adv Med Educ Pract”[journal] OR “BMC Med Educ”[journal] OR “Can Med Educ J”[journal] OR “Clin Teach”[journal] OR “J Contin Educ Health Prof”[journal] OR “Teach Learn Med”[journal] OR “Perspect Med Educ”[journal] OR “Med Educ”[journal] OR “Med Educ Online”[journal] OR “Med Teach”[journal] OR “J Grad Med Educ”[journal] OR “Int J Med Educ” [journal])

### Web of Science (LIMIT: 1999/01/01 – 2019/12/31)

((TI=(“knowledge synthesis”) OR AB=(“knowledge synthesis”)) OR (TI=(“evidence synthesis”) OR AB=(“evidence synthesis”)) OR (TI=(“literature review”) OR AB=(“literature review”)) OR (TI=(“evidence synthesis”) OR AB=(“evidence synthesis”)) OR (TI=(“systematic review”) OR AB=(“systematic review”) OR (TI=“(review) OR AB=(review)) OR (DT=(review)) OR (TI=(scoping) OR AB=(scoping)) OR (TI=(“meta-synthesis”) OR AB=(“meta-synthesis”)) OR (TI=(“narrative review”) OR AB=(“narrative review”)) OR (TI=(“critical review”) OR AB=(“critical review”)) OR (TI=(“critical synthesis”) OR AB=(““critical synthesis”)) OR (TI=(“integrative review”) OR AB=(“integrative review”)) OR (TI=(“integrative synthesis”) OR AB=(“integrative synthesis”)) OR (TI=(“qualitative review”) OR AB=(“qualitative review”)) OR (TI=(“metastudy”) OR AB=(“metastudy”)) OR (TI=(“realist review”) OR AB=(“realist review”)) OR (TI=(“critical review”) OR AB=(“critical review”)) OR (TI=(“critical review”) OR AB=(“critical review”)) OR (TI=(“rapid review”) OR AB=(“rapid review”)) OR (TI=(“umbrella review”) OR AB=(“umbrella review”)) OR (TI=(“BEME”) OR AB=(“BEME”)) OR (TI=(“consensus conference”) OR AB=(“consensus conference”)) OR (TI=(“medline”) OR AB=(“medline”)) OR (TI=(“cinahl”) OR AB=(“cinahl”)) OR (TI=(“pubmed”) OR AB=(“pubmed”)) OR (TI=(“embase”) OR AB=(“embase”)) OR (TI=(“psychinfo”) OR AB=(“psychinfo”))) AND SO=((Advances in Health Sciences Education) OR (Clinical Teacher) OR (Teaching “and” Learning in Medicine) OR (Medical Education Online) OR (Medical Teacher))

